# α-synuclein in the retina leads to degeneration of dopamine amacrine cells impairing vision

**DOI:** 10.1101/760603

**Authors:** Marrocco Elena, Esposito Federica, Tarallo Valeria, Carboncino Anna, Alvino Filomena Grazia, De Falco Sandro, Franco Brunella, De Risi Maria, Alessia Indrieri, Surace Enrico Maria, De Leonibus Elvira

## Abstract

**Introduction:** α-synuclein aggregates have been identified in the retina of Parkinson’s disease patients associated to vision impairment. In this study, we sought to determine the effects of α-synuclein overexpression on the survival and function of dopaminergic (DA) amacrine cells in the retina.

**Methods:** Adult mice were intravitreally injected with an adeno-associated viral (AAV) vector to overexpress human wild-type α-synuclein in the inner retina. Following systemic injections of levodopa (L-DOPA), retinal responses and visual acuity driven behavior were measured by electroretinography (ERG) and water maze task, respectively. Amacrine cells and ganglion cells were counted at 1, 2 and 3 months post-injection.

**Results:** α-synuclein led to an early loss of DA cells, which was associated with the decrease of light-adapted ERG responses and visual acuity. Systemic injections of L-DOPA rescued these retinal and visual abnormalities.

**Conclusions:** The data show that α-synuclein affects dopamine neurons in the retina. The approach provides a novel accessible mode of modeling the underlying mechanisms implicated in synucleinophaties pathogenesis and for testing novel treatments.

## INTRODUCTION

The two hallmarks of Parkinson’s disease (PD) are the formation of α-synuclein (α-syn) inclusions into Lewy bodies (LBs)^1^ and the degeneration of the nigrostriatal dopaminergic systems leading to motor and cognitive deficits in PD. Experimental evidence in animal models established a causal link between α-syn overexpression or mutations and the degeneration of dopaminergic neurons, by showing that recombinant adeno-associated viral vector (rAAV)-mediated overexpression of human wild-type (WT) α-syn in the midbrain of adult rodents leads to progressive loss of nigral dopaminergic neurons^2,3^. Starting from the clinical observation that PD patients also present with visual symptoms such as reduced visual acuity, low contrast sensitivity, altered electroretinogram (ERG) and disturbed color vision^4-7^, recent findings are considering the retina as a potential biomarker for PD.

α-syn aggregates have been identified in the retina of PD patients and in particular in the Ganglion Cell Layer (GCL), Inner Plexiform Layer (IPL) and Inner Nuclear Layer (INL)^8^. According to clinical studies, transgenic mouse models overexpressing α-syn reported an accumulation of the protein in specific retinal cells depending on the promoter used for the expression^9^. A transgenic mouse model overexpressing a fused α-syn-GFP under the Platelet Derived Endothelial Growth Factor (PDGFβ) promoter reported dot-like deposits in the INL and in the retinal ganglion cells (RGC) increasing over time^10^, and leading to their degeneration. However, none of these studies explored the effects of synucleinopathy on dopaminergic (DA) amacrine cells of the retina, which are located in the INL^11-13^.

In the retina, DA neurotransmission is regulated by a single subpopulation of amacrine cells (A18 cells)^14^ that synthesizes and releases DA neurotransmitter^15,16^. DA plays a crucial role in the modulation of light adaptation. Light stimuli, through rods, cones, and melanopsin ganglion cells activate dopaminergic amacrine cells, triggering dopamine release^17,18^. Extracellular DA acts on dopaminergic receptors. In particular, D1-like receptors are expressed not only by amacrine cells but also in horizontal cells and bipolar cells; rods and cone mainly express D4R; D4R stimulation reduces the rod-cone communication through the gap junctions, which increases the direct response of cones in photopic conditions^14-16^. On the contrary, the absence of DA favors the rod-cone conductance and the shunting cone electrical signal. Retinal dopaminergic tone is thought to amplify the cone circuits flow, producing a shift from rod-dominant to cone-dominant vision during daylight^19-21^. DA cells also contribute to color vision. These visual functions are frequently compromised in Parkinsonian patients^15,22^.

In this study we show, for the first time, that α-syn overexpression in the retina leads to neurodegeneration of DA amacrine cells, causing retinal-specific defects and a consequent visual function impairment.

## MATERIAL AND METHODS

### Subjects

All the experiments were performed in male and female inbred C57BL/6J adult (12-20 weeks old) mice. Group housed mice, with *ad libitum* access to water and food, were maintained at 22±1 °C and 55±5% relative humidity, with a 12-h light: 12-h dark cycle (lights on 07:30-19:30) and tested during the light phase. The experiments were conducted in accordance with the European Communities Council directives and Italian laws on animal care.

### Intravitreal injections

Animals were intravitreally injected with the same recombinant adeno-associated viral vector (rAAV) 2/6 expressing human (hu) α-synuclein (rAAV2/6-hu-α-syn) or with a rAAV2/6 expressing GFP (rAAV2/6-GFP) (7.7 × 10^13^ genome copies/ml) previously described^2^. Intravitreal injection of rAAV2/6 has been previously reported to transduce the retina in mice and rats^23,24^. Pupils of anaesthetized animals (100 mg/kg medetomidine and 0.25 mg/kg ketamine) were dilated using 1% tropicamide and 2.5% phenylephrine (Chauvin, Essex, UK) and a small guide hole was made under the limbus with a 30G needle. The eye was gently massaged with a cotton swab to remove a portion of the vitreous to avoid a post-injection reflux of vitreous and/or drug solution. Then, 1 µL of vector was intravitreal injected through the initial hole using a 34G Hamilton syringe.

### Visual acuity test

We used a behavioral procedure of the visual acuity task modified from Prusky’s^25^ and Robison’s^26^ procedures and described in supplementary materials.

#### Electrophysiological recordings

Mice were anesthetized and accommodated in a stereotaxic apparatus; their pupils were dilated with 1% tropicamide and 2.5% phenylephrine (Chauvin, Essex, UK).

#### Dark-adapted ERG

Mice were dark-adapted for 3 hours. After anaesthesia and mydriasis, retinal electrical response was recorded after flashes of different light intensities ranging from – 4 to + 1 log cd.s/m2, in scotopic condition. For each animal group, mean of A-wave and B-wave amplitudes were plotted in relation to the intensity of light that evoked them.

#### Light-adapted ERG

Retinal electrical response was recorded in the presence of a constant background illumination set at 50 cd/m2 and after 10-ms flashes (white light) of 10.0 cd s/m2 elicited at different time intervals (from 0 minute to 8 minute; to intervals 2 minute for each step)^20^. Flashes were generated through a Ganzfeld stimulator (CSO, Florence, Italy). For each animal group, the mean of B-wave recorded over time was plotted.

### Drugs

Benserazide (20 mg/kg, Sigma Aldrich) in PBS 1x was injected intraperitoneally (i.p.) injected 15 minutes before L-DOPA (10 mg/kg, Sigma Aldrich, in saline solution, i.p.) or saline. Each animal received L-DOPA and saline 5-7 days apart in randomized order.

### Immunofluorescence

One, two or three months after rAAV-injection, the eyes were removed and fixed in 4% paraformaldehyde fixative solution (PFA), cryoprotected in PBS sucrose, embedded in OCT (Optimal cutting temperature) and cut by cryostat at 12 μm. Sections were processed by immunofluorescence as previously described^2^, using the following primary antibodies: TH (AB 152 Merck Millipore), hu-α-syn (AB 211 Santa Cruz Biotechnology), hu-α-syn phosphorylated at Ser129 (AB 51253, Abcam), GAD65 (198 102 Synaptic System), NeuN (AB 104225 Abcam), RBPMS (15187-AP Proteintech), ChAT (AB144P Chemicon) diluted 1: 400. DAPI as a nuclear counterstain was used. Images were taken using a fully motorized Nikon microscope ECLIPSE Ni-E. Quantification of cells number was performed by direct counting at microscope on post-immunofluorescence vertical retinal sections in the INL for GAD65, TH and ChAT and in the GCL for NeuN and RBPMS. For each eye, an average of 18 serial retinal sections of 12 µm tick each was counted. For TH staining, retina flat mounts are obtained by three radial cuts from optic nerve. The counting is performed by ImageJ software; for each retina sample, three representative areas (at a magnification of 10X) on flat mount were quantized.

## Results

### α-synuclein overexpression in the retina leads to vision impairment, which is rescued by L-DOPA administration

α-syn was expressed in the retina of adult C57BL/6J mice, through an intravitreal bilateral injection of rAAV-hu-α-syn (experimental cohort) or rAAV-GFP (control cohort). We used a within subject experimental design, by testing the animal abilities to adapt to light changes under rod saturating conditions with ERG and assess their visual acuity in the water maze task, before and after rAAV injection (Fig. **1A**). To mimic the adaptation to light, a series of flashes at a distance of 2 min are evoked in the presence of a constant light background. The B-wave amplitude (more detectable in rodents in photopic condition^27^ than A-wave) will be considered as index of retinal response following light adaptation ERG analysis; rAAV-hu-α-syn injected mice showed a selective impairment of light-adapted responses (time dependent decrease; Fig. **1B**), despite conservation of those elicited in dark-adapted conditions (Fig. **1C, D**). The amplitude defect of B-wave in light-adapted condition (Fig. 1B) was completely rescued by DA replacement via L-DOPA systemic administration (Fig. **1G**). These ERG findings are consistent with those performed in mice with retinal specific genetic ablation of tyrosine hydroxylase (TH), the key enzyme of the DA biosynthetic pathway^20^.

**Figure 1.**
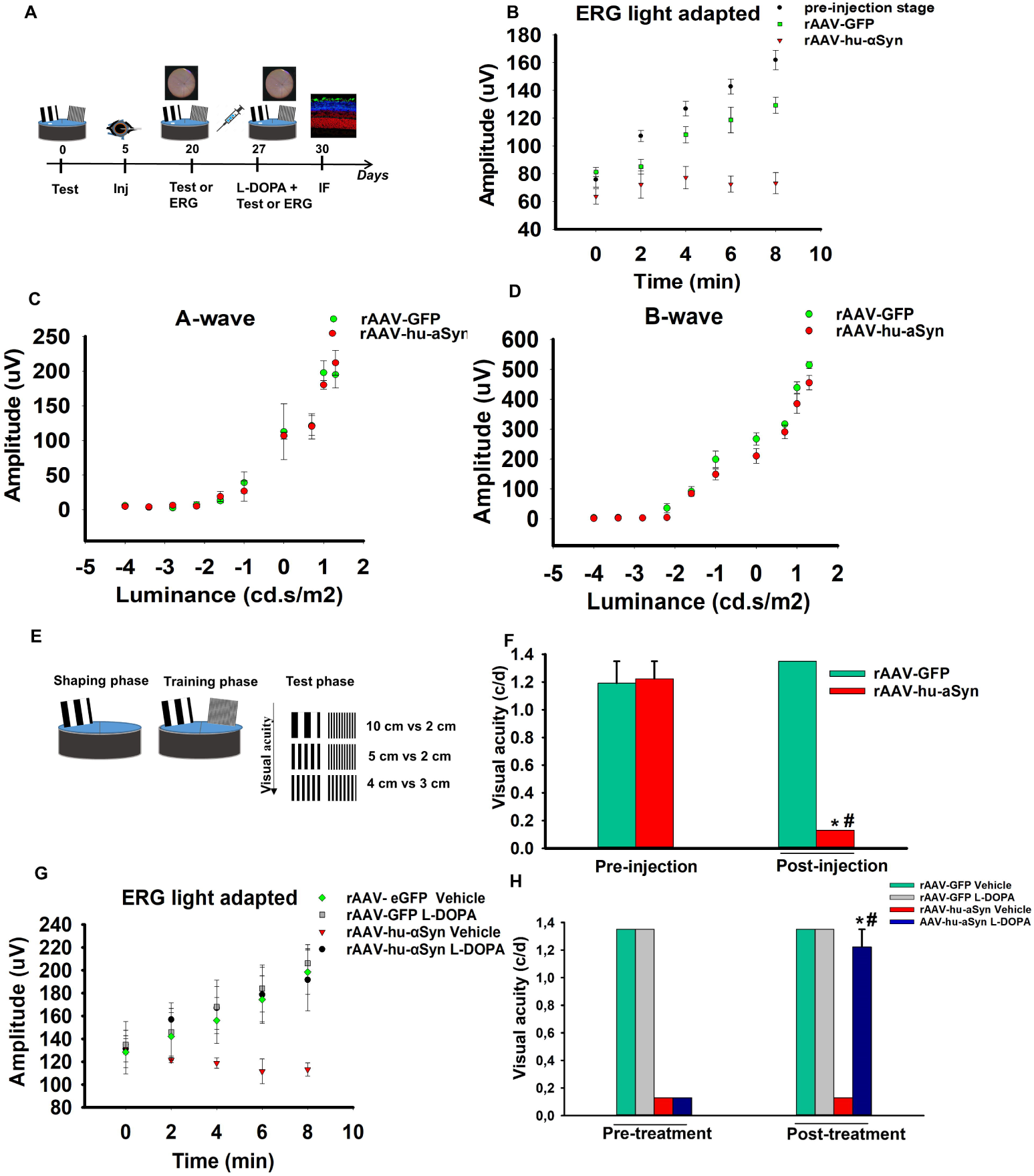
Visual function impairment in rAAV-hu-α-syn injected mice are rescued by L-DOPA. (**A**) Experimental design reporting the temporal order (days) of the experiments: the visual acuity test or ERG analysis before and after the rAAV injection (inj) and after L-DOPA or vehicle treatment followed by finally the immunofluorescence analysis (IF). (**B**) Light-adapted ERG measured over time (min), shows a severe and selective decrease of B-wave amplitude (mean) in rAAV-hu-α-syn injected mice compared to both rAAV-GFP mice and to the pre-injection time point [repeated measures ANOVA group F _(2,16)_ = 16.428; p = 0.0001; time F _(4,64)_ = 30.673; p < 0.0001, groups x time F _(8,64)_ = 11.473; p < 0.0001; (pre-injection, *n = 10*; rAAV-GFP, *n = 4*; rAAV-hu-α-syn *n = 5*), Duncan *post hoc* test]. (**C-D**) No differences in A-wave and B-wave (mean) measured in the ERG dark-adapted condition were observed between rAAV-hu-α-syn and rAAV-GFP mice [**C:** repeated measure ANOVA group F_(1,9)_ = 0.018; p = 0.8951; luminance F_(9,81)_ = 88.406; p < 0.0001; group x luminance F_(9,81)_ = 0.321; p = 0.9659; (rAAV-GFP, *n = 5*; rAAV-hu-α-syn, *n = 6*); **D:** repeated measure ANOVA group F(1,10) = 5.763; p = 00373; luminance F_(9,90)_ = 217.661; p < 0.0001; group x luminance F_(9,90)_ = 1.094; p = 0.3755; (rAAV-GFP, *n = 5*; rAAV-hu-α-syn, *n = 8*)]. (**E**) Schematic representation of the different phases of the visual acuity test: 10 cm vs 2, 5 cm vs 2 cm, 4 cm vs 3 cm indicate the couples of cards used. (**F**) Impaired visual acuity (cycles per degrees, c/d) in rAAV-hu-α-syn inejcted mice compared to the pre-injection stage and rAAV-GFP mice [repeated measures two-way ANOVA, group F_(1,7)_ = 35.028; p = 0.0006; stage F_(1,7)_ = 21.635; p = 0.0023; group × stage F_(1,7)_ = 38.878; p = 0.0004; (rAAV-GFP, *n = 4*; rAAV-hu-α-syn, *n = 5*) Duncan *post hoc* test]. (**G**) L-DOPA rescued the light-adapted ERG defect in rAAV-hu-α-syn [repeated measures ANOVA, group × treatment × time F_(4,72)_ = 2.201; p = 0.0774; (rAAV-GFP Vehicle, *n = 4*; rAAV-GFP L-DOPA, *n = 4*; rAAV-hu-α-syn Vehicle, *n = 4*; rAAV-hu-α-syn L-DOPA, *n = 10*)]. (**H**) L-DOPA rescued the visual acuity defect in rAAV-hu-α-syn injected mice [repeated measures three-way ANOVA, rAAV group x treatment x stage F _(1,14)_ = 57.407; p < 0.0001; (rAAV-GFP Vehicle, *n = 4*; rAAV-GFP L-DOPA, *n = 4*; rAAV-hu-α-syn Vehicle, *n = 5*; rAAV-hu-α-syn L-DOPA, *n = 5*) Duncan *post hoc* test]. Data represent mean ± SEM. # p < 0.05 vs rAAV-GFP injection; * p < 0.05 vs rAAV-hu-α-syn vehicle.

In order to evaluate whether α-syn overexpression in the retina leads to changes in visual acuity, we investigated the ability of α-syn injected mice to discriminate between two different spatial frequencies in the water maze task using behavioral procedures similar to those previously published (Fig. **1E**, see Materials and Methods)^28^. Untreated C57BL/6J mice showed maximal visual acuity of about 1.3 c/d (Supplementary Video 1A). rAAV-hu-α-syn injection significantly impaired visual acuity (0.130 c/d) compared to both the pre-injected and rAAV-GFP injected mice (Fig. **1F**, Supplementary Video 1B). A major advantage of this intravitreal selective expression of α-syn is that it allows to specifically relate the behavioral defects in the water maze with vision impairment, ruling out the role of motor impairments characterizing genetic animal models of synucleinopathy. Systemic administration of L-DOPA rescued the visual acuity defect to pre-treatment condition (Fig. **1H**) (Supplementary Video 1C). The ameliorative effects of L-DOPA were specific to rAAV-hu-α-syn injected eyes, as L-DOPA did not affect rAAV-GFP injected mice and CD-1 outbred albino mice (Supplementary results and Fig. **1A-B**).

Thus, the selective complementation results obtained by administering L-DOPA in α-syn injected mice support that the visual defect induced by α-syn overexpression are specifically due to the loss of DA amacrine neurotransmission. These results parallel those obtained after selective DA neuronal loss induced by intravitreal administration of the DA selective neurotoxin, 6-hydroxydopamine (6-OHDA), resulting in a significant and specific ablation of the TH+ amacrine cells, leading to a visual-acuity impairment similarly rescued by L-DOPA supplementation (Supplementary Fig. **2A-D**).

### α-synuclein overexpression leads to a time-dependent dopamine neuronal loss

To determine whether the underlying mechanisms of the functional findings correspond to specific histopathological effects we analyzed by immunofluorescence the rAAV injected retinas. At one-month post intravitreal administration, rAAV2/6-GFP transduced efficiently the inner retina including IPL, the deepest INL and extensively the GCL (Fig. **2A**). In addition, rAAV-hu-α-syn injected retinas showed also positive to phosphorylated α-syn, at serine 129, a specific biomarker of α-syn aggregates and/or misfolded protein^8,29^. Accordingly, to the functional data, we found a highly significant and progressive (1, 2 and 3 months) reduction of the number of TH-immunoreactive amacrine cells in the retinas, evaluated with both vertical and whole mount slices, of rAAV-hu-α-syn injected mice (Fig. **2B-C**). Notably, we found an unchanged number of GABAergic glutamic acid decarboxylase 65 (GAD65)-positive amacrine cells and cholinergic amacrine cells expressing Choline acetyltransferase (ChAT) at this early stage (Supplementary Fig. **3C-D**). Ganglion cell numbers was not affected at one month after the injection, as evaluated through NeuN staining. However, a significant reduction of NeuN+ cells number, was detectable at a later time point (3 months), albeit not to the same extent as that observed in DA amacrine cells (Fig. **2D-E**). As NeuN in the ganglion cells layer marks both ganglion cells and displaced amacrine cells, we performed RBPMS staining that is selectively expressed in adult ganglion cells, to confirm a degeneration of this neuronal population at this later stage (3 months) (Fig. **2F-G**).

## Discussion

The observation in Parkinsonian patients of a reduced DA innervation in the central retina associated with altered adaptation to light contrast and specific defects in visual acuity was made already 30 years ago^30^. Experimental evidence also provided in genetic and pharmacological animal models, in which loss of function of TH+ amacrine cells results in visual defect that can be corrected by L-DOPA replacement therapy^20,31-33^. More recent evidence suggested that α-syn aggregates, are present in the eye of PD patients, as also confirmed by transgenic animal models of α-synucleinopathy^9,10^, and that phosphorylated α-syn accumulates in the retina in parallel with that in the brain, including in early stages preceding development of clinical signs of parkinsonism or dementia^34^. However, synucleinopathy in the retina has never been directly associated to DA cells neurodegeneration and consequent with visual defects.

In this study, we report for the first time that α-syn overexpression in the retina induces a time-dependent loss of TH+ amacrine cells which precedes ganglion cell degeneration. In this model, no detectable changes in the number of GABAergic and cholinergic amacrine cells were observed. Many of the TH+ positive cells co-express GABA; however, TH+ are only a minor part of GABAergic amacrine cells in the INL. Therefore the lack of significant changes in the number of GABA positive amacrine cells might be explained by considering that DA and GABAergic are only a minimal part of the latter, respectively^11-13^.

The loss of TH+ cells was associated to altered light-adaptation and impaired performance in the visual acuity version of the water maze task.

Although, the histological, electrophysiological and behavioral findings we provide in this study consistently suggest that the impairment in the early stage was selectively induced by DA amacrine cells function, we cannot completely exclude that other neuronal population might be affected at this early stage. Nevertheless, the fact that systemic injection of L-DOPA, which selectively acts on DA neurons, completely rescued the visual defects strongly suggests that the early functional impairments induced by α-syn overexpression are selectively dependent on impaired DA cellular function. These findings show that α-syn exerts harmful effects on DA neurons independently from the cellular context in which they are integrated, including the retina.

The eye is gaining momentum for studying neurodegenerative disorders due to its easy accessibility^35^. Interestingly, we could detect phosphorylated α-syn at this early stage in the retina, which has been suggested as a valid early marker of the pathology^34^. Indeed, it is being considered as an ideal model organ for both an early identification of protein aggregates and for testing novel therapeutic approaches in neurodegenerative disorders.

## Acknowledgements

We thank Prof. Anders Bjorklund and Jenny G. Johansson for providing the vector. We thank Prof. Alberto Auricchio and Prof. Barbara Picconi for critically revising the manuscript. We also thank De Leonibus’ lab for critical revision of the manuscript. We thank Martina Colucci for helping with histological experiments, and TIGEM’ Animal House and Microscopy Facilities.

## Supplementary materials and methods

### Visual acuity test

#### Apparatus

Animals were tested in a circular tank (150 cm diameter, 35.5 cm high) filled with water (22±1 °C). A rectangular white platform (37 cm long × 13 cm wide × 14 cm high) was submerged 1 cm below the water surface and a steel divider (70 cm long) extending toward the center of the pool, divided it in two equal quadrants; the divider constituted the response choice point. Two cards (40×44.5 cm) with vertical pattern black and white stripes of different width were fixed to the wall of the pool in each quadrant. The escape platform was located in front of the card with smaller stripes. Animals discriminated between a card with larger stripes and a card with smaller stripes where the escape platform was located. Correct responses were recorded as direct entry in the quadrant where the card with smaller stripes was located. Visual acuity was finally measured in cycles/degree (c/d) according to the method described in^28^.

#### Procedure

The task consisted of three phases: 1. During the shaping phase (day1), animals were habituated to the task, by positioning the card with 10 cm black and white stripes and the platform in the same quadrant so they learned that a platform was associated to a card. The platform and the card were alternatively positioned in left or right quadrant for 3 sessions of 6 trials. Animals were released in the pool at progressively greater distance from the choice point in each session. 2. Training phase: the 10 cm black and white striped card and the 1 cm black and white striped card were randomly positioned in left or right quadrant and the platform was always located under the smaller striped card. Animals were trained to discriminate between two cards and to learn that the platform was always associated to smaller striped card. They were tested for a maximum of 3 sessions per day (10 trials/session, 60 seconds/trial). If the animal made 70% correct responses (criterion) in a session or 4 consecutive correct responses, it was tested in the next phase; if the animals did not reach the criterion, they were re-tested in the training phase for a maximum of 3 days (3 sessions per day) 3. Test phase: couples of cards used were progressively more difficult to discriminate: 10 cm striped card vs. 1 cm striped card, 10 cm striped card vs. 2 cm striped card, 5 cm striped card vs 2 cm striped card, 4 cm striped card vs 3 cm striped card. To go to the next couple of cards mice were required to make 70% corrects responses or 4 consecutive correct responses at the previous couple of cards. Animals were tested for a maximum of 3 sessions per day (10 trials/session, 60 seconds/trial).

### 6-hydroxidopamine experiment

A group of mice was intravitreally injected with 6-OHDA (2µg/µL, Sigma Aldrich, 1µL/side) and a group was injected with saline as control. Intravitreal injections were performed as described for viral vector injection (see Materials and Methods). 15-20 days after the injection procedure, mice underwent the behavioral procedure. Systemic L-DOPA or vehicle treatment started 1 week after the injection. The protocol used was the same described for the rAAV injected mice. Each animal was first administered with L-DOPA (n=7) or Vehicle (n=5), one week apart with the inverse treatment (Supplementary Fig. **2A-B**).

Male and female outbred CD-1 mice (12-20 weeks old) (n=10) were also used to validate our behavioral procedure. 6 of them were treated with L-DOPA and 4 of them were treated with vehicle.

### Statistical analysis

Statistical analysis was performed using a two-way ANOVA for repeated measures for the behavioral analysis before the L-DOPA treatment (between variable: Control and 6-OHDA; repeated measures: pre and post-lesion) and a three-way ANOVA for repeated measures for the behavioral analysis under L-DOPA treatment (between variables, Group: Control or 6-OHDA, treatment: L-DOPA and Vehicle; repeated measures: pre and post-treatment). TH+ Cells counting was analysed with a one-way ANOVA. Behavioral experiments in CD1 mice were analysed with a one-way ANOVA (between variable: CD-1 and C57BL/6J) or with a two-way ANOVA for the experiment with the L-DOPA treatment. Duncan *post-hoc* test was used when appropriate and the statistical significance was set at p < 0.05.

## Supplementary Results

### 6-OHDA injection caused similar visual acuity defects to those observed in rAAV-hu-α-syn injected mice, rescued by L-DOPA treatment

Intravitreal injection of 6-OHDA in adult mice caused a 50% reduction of TH+ amacrine cells 1 month post-injection (Supplementary Fig. 2A-B). 15 days after the injection, 6-OHDA injected mice showed a significant impairment in the visual acuity task so that they did not overcome the first step of the task (10 cm vs 1 cm, 0.313 c/d) (Supplementary Fig 1C). L-DOPA treatment completely rescued the defect (Supplementary Fig. 1D).

**Figure 2.**
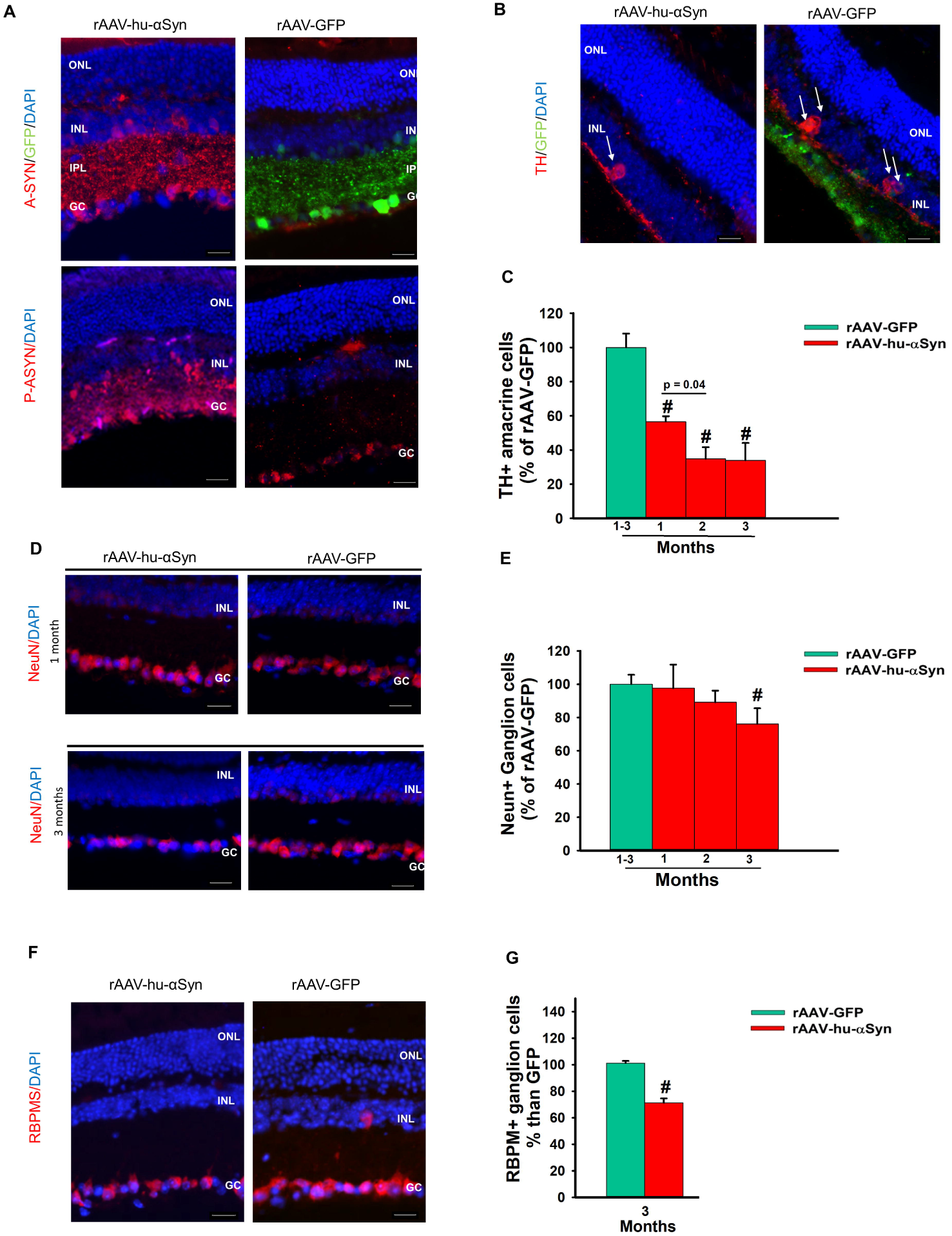
Time-dependent reduction of TH+ amacrine cells number in rAAV-hu-α-syn injected mice. (**A**) Representative immunofluorescence showing retinal sections of rAAV-hu-α-syn injected mice and rAAV-GFP mice stained with antibody anti-hu-α-syn (A-SYN) (red) and antibody anti-phospho α-syn (P-ASYN) (red) (20x Magnification, Scale bar 50µm). (**B**) Representative immunofluorescence on retinal sections of rAAV-hu-α-syn injected mice (left) and rAAV-GFP mice (right) stained with antibody anti-tyrosine hydroxylase (TH) (red). White arrows indicate TH+ amacrine cells 1 month post-injection (10x Magnification, Scale bar 50µm). (**C**) TH+ amacrine cells number is reduced in rAAV-hu-α-syn injected mice already at 1 month. A significant reduction was observed at 2 months compared to 1 month in rAAV-hu-α-syn groups (p =0.0419). (**D**) Representative immunofluorescence showing retinal sections of rAAV-hu-α-syn injected mice (left) and rAAV-GFP mice (right) stained with antibody anti-NeuN -(red) at 1 and 3 months post injection (20x Magnification, Scale bar 50µm). **(E)** Reduction of NeuN+ amacrine cells at 3 months post injection in rAAV hu α syn injectd mice than rAAV GFP mice. (**F**) RBPMS staining (red) performed at 3 months post injection in rAAV-hu-α-syn injected mice (left) and rAAV-GFP mice (right) (20x Magnification, Scale bar 50µm). (**G**) Number of ganglion cells RBPMS+ significantly reduced at 3 months post-injection in rAAV-hu-α-syn injected mice compared to rAAV-GFP mice [TH: one-way ANOVA, group F _(3,19)_ = 19.842; p<0.0001; (1-3 months: rAAV-GFP *n = 7*; rAAV-hu-α-syn *n = 6; 6 and 4 at 1, 2 and 3 months respectively*; Duncan *post-hoc* test; NeuN: one-way ANOVA, group (rAAV-GFP vs rAAV-hu-α-syn 3 months F _(1,27)_ = 5.204; p = 0.0306; (1-3 moths: rAAV-GFP, (n *= 20*); rAAV-hu-α-syn, *n = 6*); Duncan *post-hoc* test; RBPMS: one-way ANOVA, group (rAAV-GFP vs rAAV-hu-α-syn 3 months F _(1,27)_ = 5.204; p = 0.0306; (1-3 moths: rAAV-GFP, (n *= 20*); rAAV-hu-α-syn, *n = 6; 6 and 4; at 1, 2 and 3 months respectively*);]. Data represent mean ± SEM. # p < 0.05 *vs* rAAV-GFP. * p < 0.05 vs. rAAV-hu-α-syn 1 month.

### L-DOPA treatment did not rescue visual acuity defect in CD-1 mice

To further assess the specificity of the effects of L-DOPA on DA, we tested CD-1 albino outbred mice, which have normal levels of DA and L-DOPA, but reduced melanin due to the lack of tyrosinase action on the melanin biosynthetic pathway^36,37^. As expected, albino mice showed reduced visual acuity relatively to pigmented mice (Supplementary Fig. **1A**)^38^ and L-DOPA treatment was not sufficient to rescue the defects, in line with previous clinical findings showing similar negative results in albino patients^39^ (Supplementary Fig. **1B**).

**Supplementary Figure 1.** (**A**) Visual acuity measured in cycles per degrees (c/d) in CD-1 and in C57BL/6J mice (C57BL/6J n = 10; CD-1 n = 10). (**B**) L-DOPA or Vehicle treatment in CD-1 mice (Vehicle n = 4; L-DOPA n = 6). * p < 0.05 vs C57BL/6J.

**Supplementary Figure 2. Reduced visual acuity in 6-OHDA injected mice is rescued by L-DOPA.** (**A**) Retinal sections showing TH positive (TH+) amacrine cells (green signal) in control and 6-OHDA injected mice (10X magnification, scale bar 50µm). (**B**) Number of TH+ amacrine cells in control and 6-OHDA injected mice [one-way ANOVA, group F _(1,9)_ = 8.687; p= 0.0163, Control n = 4; 6-OHDA n = 7; Duncan post-hoc test]. (**C**) Visual acuity measured in cycles per degrees (c/d) in the pre-lesion and in the post-lesion stage. 6-OHDA caused a significant impairment in visual acuity [repeated measures ANOVA, group F_(1,22)_ = 46.720; p < 0.0001, stage F_(1,22)_ = 481.467; p < 0.0001, groups x stage F_(1,22)_ = 481.467; p < 0.0001; Control *n = 11*; 6-OHDA *n = 13*; Duncan *post hoc* test]. (**D**) L-DOPA rescue in 6-OHDA injected mice (Control vehicle *n = 3*; Control L-DOPA *n = 3*; 6-OHDA *n = 9*; 6-OHDA *n = 9*). Data represent mean ±SEM. # p < 0.05 vs Control, vs 6-OHDA Vehicle; * p < 0.05 vs pre-lesion, pre-treatment stage.

**Supplementary Figure 3. Dopaminergic, but not GABAergic or cholinergic amacrine cells, are lost in α-syn injected retina at early time point.** (**A**) Representative micrographs from three quadrants of the retina flat mount (shown in right scheme) of TH staining at 1 month post injection of rAAV-hu-α-syn compared to the control retinas injected with rAAV-GFP. (**B**) Dopaminergic amacrine cells count on TH retina flat mount staining, after injection of rAAV-hu-α-syn; results are reported in percentage (%) from retina injected with GFP [one-way ANOVA, group F_(1,3)_=1161.600; p < 0.0001, rAAV-hu-α-syn *n=3*; rAAV-GFP *n=2*] (**C**) GAD65 and ChAT staining on retina of rAAV-hu-α-syn and rAAV-GFP-injected mice. (**D**) ChAT+ and GAD65 cells count at 1 month post injection [ChAT+: one-way ANOVA, group F (1,28) = 2.128; p = 0.155, rAAV-hu-α-syn *n=2*; rAAV-GFP *n=2*; GAD65: one-way ANOVA, group F (1,41) = 0.733; p = 0.3970, rAAV-hu-α-syn *n=2*; rAAV-GFP *n=2*].

**Supplementary Videos.** Videos showing visual task performance of C57BL/6J mice pre-injection (Video 1), after intravitreal injection of (rAAV-hu-α-syn) combined with saline (Video 2) and L-DOPA replacement therapy (Video 3). (Video 1) Representative sample trial of a C57BL/6J mice during the visual task requiring where to discriminate a 3 cm black and white striped card from a 4 cm black and white striped card. The platform is located under the 3 cm striped card. (Video 2) Representative trial of the visual task showing that bilateral rAAV-hu-α-syn intravitreal injection affects performance at the first stage of the procedure when animals are required to discriminate a 1 cm black and white striped card from a 10 cm black and white striped card. The platform is located under the 1 cm striped card. (Video 3) Representative trial of the visual task showing that L-DOPA treatment rescues performance in rAAV-hu-α-syn injected mice on the same trial reported in the Video 2.

## References

1. Spillantini MG, Schmidt ML, Lee VM, Trojanowski JQ, Jakes R, Goedert M. Alpha-synuclein in Lewy bodies. Nature. Aug 28 1997;388(6645):839–840.

2. Giordano N, Iemolo A, Mancini M, et al. Motor learning and metaplasticity in striatal neurons: relevance for Parkinson’s disease. Brain. Feb 1 2018;141(2):505–520.

3. Lundblad M, Decressac M, Mattsson B, Bjorklund A. Impaired neurotransmission caused by overexpression of alpha-synuclein in nigral dopamine neurons. Proc Natl Acad Sci U S A. Feb 28 2012;109(9):3213–3219.

4. Skrandies W, Gottlob I. Alterations of visual contrast sensitivity in Parkinson’s disease. Hum Neurobiol. 1986;5(4):255–259.

5. Bodis-Wollner I, Marx MS, Mitra S, Bobak P, Mylin L, Yahr M. Visual dysfunction in Parkinson’s disease. Loss in spatiotemporal contrast sensitivity. Brain. Dec 1987;110 (Pt 6):1675–1698.

6. Jones RD, Donaldson IM, Timmings PL. Impairment of high-contrast visual acuity in Parkinson’s disease. Mov Disord. 1992;7(3):232–238.

7. Matsui H, Udaka F, Tamura A, et al. Impaired visual acuity as a risk factor for visual hallucinations in Parkinson’s disease. Journal of geriatric psychiatry and neurology. Mar 2006;19(1):36–40.

8. Beach TG, Carew J, Serrano G, et al. Phosphorylated alpha-synuclein-immunoreactive retinal neuronal elements in Parkinson’s disease subjects. Neuroscience letters. Jun 13 2014;571:34–38.

9. Surguchov A, McMahan B, Masliah E, Surgucheva I. Synucleins in ocular tissues. Journal of neuroscience research. Jul 1 2001;65(1):68–77.

10. Price DL, Rockenstein E, Mante M, et al. Longitudinal live imaging of retinal alpha-synuclein::GFP deposits in a transgenic mouse model of Parkinson’s Disease/Dementia with Lewy Bodies. Scientific reports. Jul 8 2016;6:29523.

11. May CA, Nakamura K, Fujiyama F, Yanagawa Y. Quantification and characterization of GABA-ergic amacrine cells in the retina of GAD67-GFP knock-in mice. Acta ophthalmologica. Jun 2008;86(4):395–400.

12. Ghinia MG, Novelli E, Sajgo S, Badea TC, Strettoi E. Brn3a and Brn3b knockout mice display unvaried retinal fine structure despite major morphological and numerical alterations of ganglion cells. The Journal of comparative neurology. Jan 1 2019;527(1):187–211.

13. Keeley PW, Reese BE. Morphology of dopaminergic amacrine cells in the mouse retina: independence from homotypic interactions. The Journal of comparative neurology. Apr 15 2010;518(8):1220–1231.

14. Kolb H. Anatomical pathways for color vision in the human retina. Visual neuroscience. Jul-Aug 1991;7(1-2):61–74.

15. Witkovsky P. Dopamine and retinal function. Documenta ophthalmologica. Advances in ophthalmology. Jan 2004;108(1):17–40.

16. Puopolo M, Hochstetler SE, Gustincich S, Wightman RM, Raviola E. Extrasynaptic release of dopamine in a retinal neuron: activity dependence and transmitter modulation. Neuron. Apr 2001;30(1):211–225.

17. Zhao X, Wong KY, Zhang DQ. Mapping physiological inputs from multiple photoreceptor systems to dopaminergic amacrine cells in the mouse retina. Scientific reports. Aug 11 2017;7(1):7920.

18. Perez-Fernandez V, Milosavljevic N, Allen AE, et al. Rod Photoreceptor Activation Alone Defines the Release of Dopamine in the Retina. Current biology: CB. Mar 4 2019;29(5):763–774 e765.

19. Ribelayga C, Mangel SC. Identification of a circadian clock-controlled neural pathway in the rabbit retina. PLoS One. Jun 10 2010;5(6):e11020.

20. Jackson CR, Ruan GX, Aseem F, et al. Retinal dopamine mediates multiple dimensions of light-adapted vision. The Journal of neuroscience: the official journal of the Society for Neuroscience. Jul 4 2012;32(27):9359–9368.

21. Kolb H. Roles of Amacrine Cells. In: Kolb H, Fernandez E, Nelson R, eds. Webvision: The Organization of the Retina and Visual System. Salt Lake City (UT)1995.

22. Armstrong RA. Visual symptoms in Parkinson’s disease. Parkinson’s disease. 2011;2011:908306.

23. Yang GS, Schmidt M, Yan Z, et al. Virus-mediated transduction of murine retina with adeno-associated virus: effects of viral capsid and genome size. Journal of virology. Aug 2002;76(15):7651–7660.

24. Hellstrom M, Ruitenberg MJ, Pollett MA, et al. Cellular tropism and transduction properties of seven adeno-associated viral vector serotypes in adult retina after intravitreal injection. Gene therapy. Apr 2009;16(4):521–532.

25. Prusky GT, West PW, Douglas RM. Behavioral assessment of visual acuity in mice and rats. Vision Res. 2000;40(16):2201–2209.

26. Robinson L, Bridge H, Riedel G. Visual discrimination learning in the water maze: a novel test for visual acuity. Behav Brain Res. Feb 15 2001;119(1):77–84.

27. Brandli A, Stone J. Using the Electroretinogram to Assess Function in the Rodent Retina and the Protective Effects of Remote Limb Ischemic Preconditioning. Journal of visualized experiments: JoVE. Jun 9 2015(100):e52658.

28. Robinson L, Harbaran D, Riedel G. Visual acuity in the water maze: sensitivity to muscarinic receptor blockade in rats and mice. Behavioural brain research. May 5 2004;151(1-2):277–286.

29. Sato H, Kato T, Arawaka S. The role of Ser129 phosphorylation of alpha-synuclein in neurodegeneration of Parkinson’s disease: a review of in vivo models. Reviews in the neurosciences. 2013;24(2):115–123.

30. J. Nguyen-Legros CH, Therese Di Paolo, Axelle Simon. The retinal dopamine system in Parkinson’s disease. Clinical Vision Sciences. 1993;8(1):1–12.

31. Citron MC, Erinoff L, Rickman DW, Brecha NC. Modification of electroretinograms in dopamine-depleted retinas. Brain research. Oct 14 1985;345(1):186–191.

32. Hempel K, Lange HW, Kayser EF, Roger L, Hennemann H, Heidland A. Role of O-methylation in the renal excretion of catecholamines in dogs. Naunyn-Schmiedeberg’s archives of pharmacology. 1973;277(4):373–386.

33. Maclin EL, Bodis-Wollner, I., and Marx, M. Simultaneous pattern eleetroretinograms and VEPs in MPTP-treated monkeys. Invest Ophthalmol Vis Sci 26:68. 1985.

34. Ortuno-Lizaran I, Beach TG, Serrano GE, Walker DG, Adler CH, Cuenca N. Phosphorylated alpha-synuclein in the retina is a biomarker of Parkinson’s disease pathology severity. Movement disorders: official journal of the Movement Disorder Society. Aug 2018;33(8):1315–1324.

35. London A, Benhar I, Schwartz M. The retina as a window to the brain-from eye research to CNS disorders. Nature reviews. Neurology. Jan 2013;9(1):44–53.

36. Lavado A, Montoliu L. New animal models to study the role of tyrosinase in normal retinal development. Frontiers in bioscience: a journal and virtual library. Jan 1 2006;11:743–752.

37. Slominski A, Zmijewski MA, Pawelek J. L-tyrosine and L-dihydroxyphenylalanine as hormone-like regulators of melanocyte functions. Pigment cell & melanoma research. Jan 2012;25(1):14–27.

38. Jeffery G. The albino retina: an abnormality that provides insight into normal retinal development. Trends in neurosciences. Apr 1997;20(4):165–169.

39. Summers CG, Connett JE, Holleschau AM, et al. Does levodopa improve vision in albinism? Results of a randomized, controlled clinical trial. Clinical & experimental ophthalmology. Nov 2014;42(8):713–721.

