## Supplementary figures and images for "α-synuclein in the retina leads to degeneration of dopamine amacrine cells impairing vision"

### Supplementary Figure 1

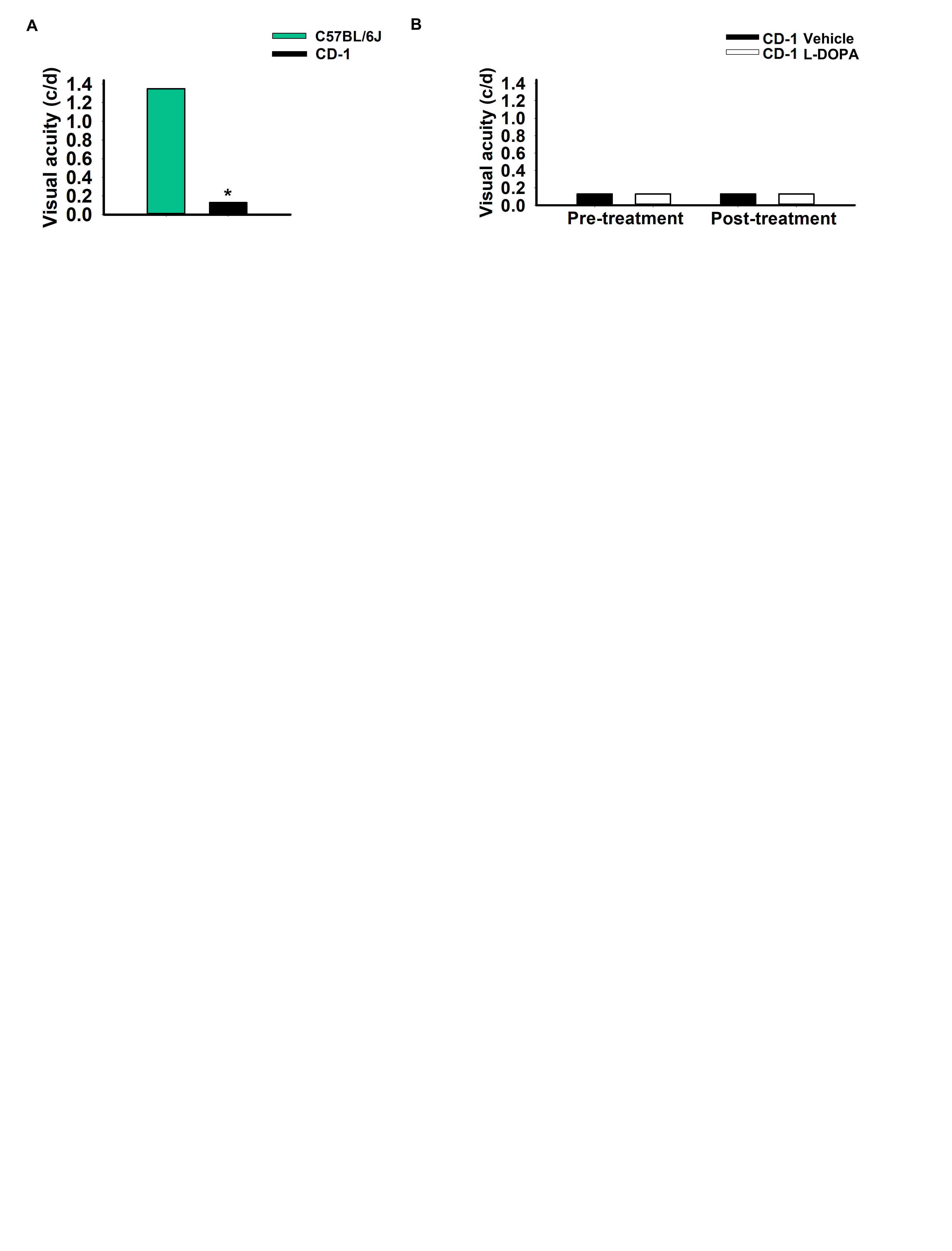

### Supplementary Figure 2

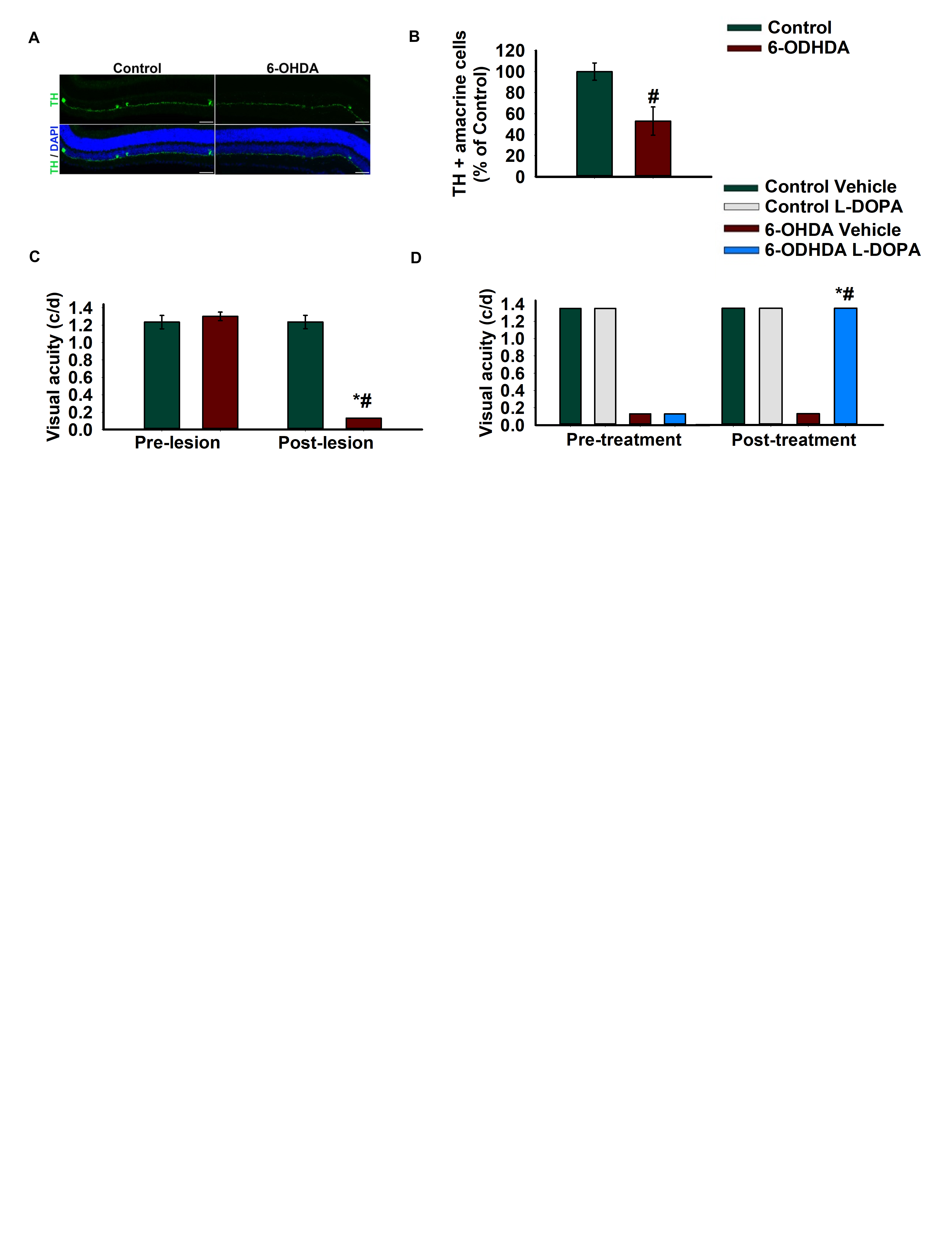

### Supplementary Figure 3

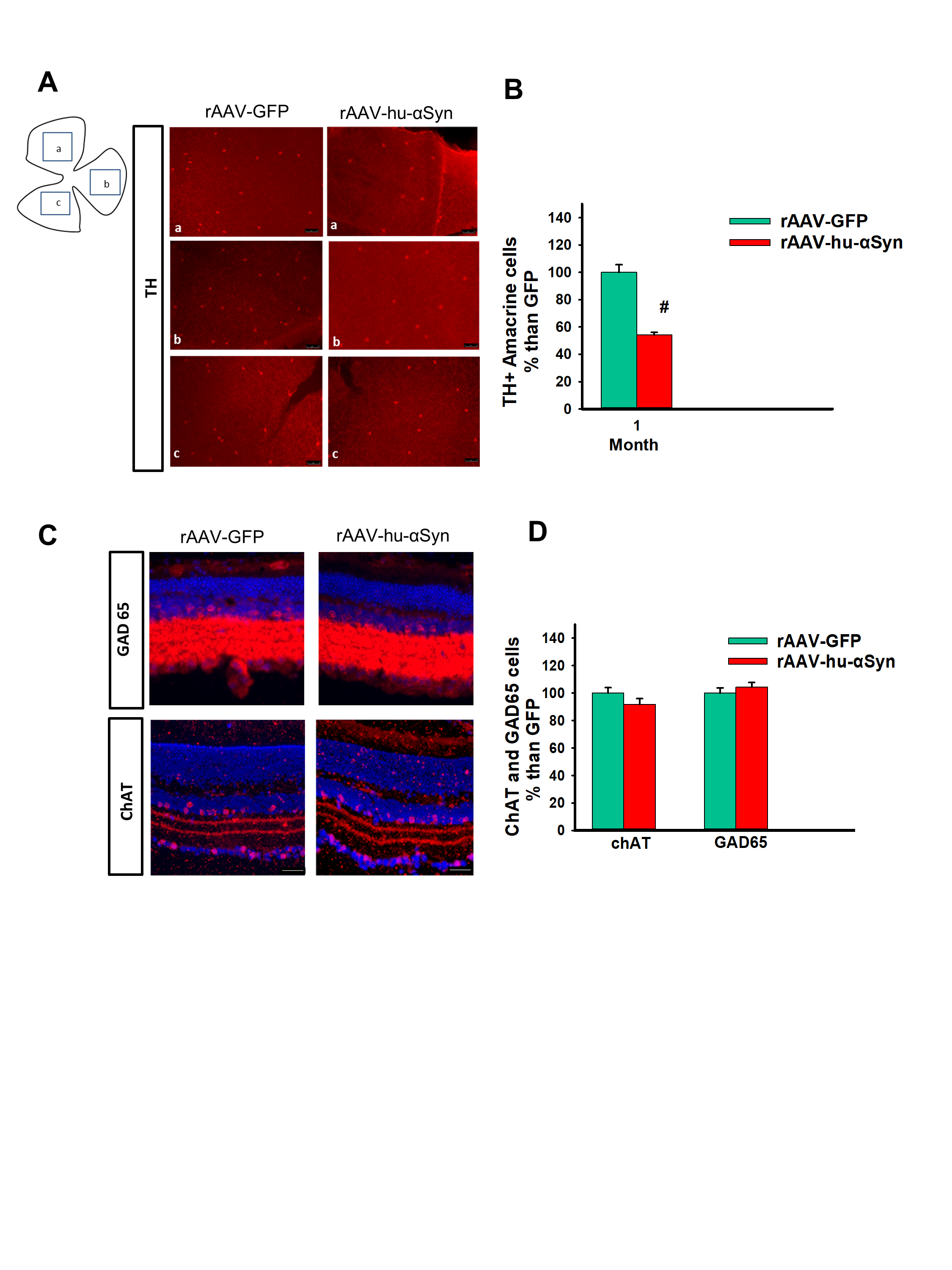
